# FRETboard: semi-supervised classification of fret traces

**DOI:** 10.1101/2020.08.28.272195

**Authors:** Carlos de Lannoy, Mike Filius, Sung Hyun Kim, Chirlmin Joo, Dick de Ridder

**Affiliations:** Bioinformatics Group, Wageningen University, 6708PB, Wageningen, The Netherlands; Department of Bionanoscience, Delft University of Technology 2629HZ, Delft, The Netherlands

**Keywords:** Förster resonance energy transfer, hidden Markov model, semi-supervised learning

## Abstract

Förster resonance energy transfer (FRET) is a useful phenomenon in biomolecular investigations, as it can be leveraged for nano-scale measurements. The optical signals produced by such experiments can be analyzed by fitting a statistical model. Several software tools exist to fit such models in an unsupervised manner, but their operating system-dependent installation requirements and lack of flexibility impede wide-spread adoption. Here we propose to fit such models more efficiently and intuitively by adopting a semi-supervised approach, in which the user interactively guides the model to fit a given dataset, and introduce FRETboard, a web tool that allows users to provide such guidance. We show that our approach is able to closely reproduce ground truth FRET statistics in a wide range of simulated single-molecule scenarios, and correctly estimate parameters for up to eleven states. On *in vitro* data we retrieve parameters identical to those obtained by laborious manual classification in a fraction of the required time. Moreover, we designed FRETboard to be easily extendable to other models, allowing it to adapt to future developments in FRET measurement and analysis.

**Availability:** source code is available at https://github.com/cvdelannoy/FRETboard. The FRETboard classification tool is also available as a browser application at https://www.bioinformatics.nl/FRETboard.

## 1 Introduction

Over the past decades, single-molecule Förster resonance energy transfer (smFRET) experiments have provided fundamental insights in biomolecular structure and many molecular mechanisms [e.g. 1–6]. Although all experiments essentially rely on the same principle – that of distance-dependent energy transfer efficiency between fluorescent donor and acceptor dyes – the use of different labeling schemes allows for versatile application. For instance, dyes fixed to two points on a single molecule may provide spatial information valuable in solving its structure [2, 4], or register structural dynamics as the molecule exerts its biological function [1]. Fixed to separate molecules, FRET may provide information on the occurrence and nature of molecular interactions [3, 5, 6]. If a single dye pair provides insufficient information, a multitude of dye pairs may even be read out simultaneously by making use of stochastically blinking dyes [7].

As such different labeling schemes produce data of a different nature, it follows that widely applicable smFRET data analysis software should be flexible enough to adapt to this variety. Several software packages exist, many of which rely on some flavor of hidden Markov model (HMM) to extract characteristic FRET efficiencies and molecular kinetics for visited molecular conformations [8–12]. Some methods only analyze the energy transfer efficiency, measured as fraction of acceptor emission intensity over total intensity [8, 10]. Others make use of the additional data contained in the individual donor and acceptor emission channels [9, 11, 12]. All aim to limit the influence of the user in setting parameters guiding the fitting process and thus adopt an unsupervised fitting procedure.

Here, we show that an HMM may be fitted to any particular FRET data set using a supervised or semi-supervised approach, i.e. by allowing the user to manually curate classification of a limited number of traces to steer model fitting (Figure 1). Such direct intervention at the classification level makes model fitting a flexible, intuitive and computationally light-weight process. We further increase accuracy by introducing a more elaborate HMM structure that, to our knowledge, has not previously been applied to smFRET data. Using several additional features derived from the original signal further boosts accuracy and increases the flexibility to adapt to data of different labeling schemes. Our method is available for use through a web tool, FRETboard. FRETboard is a start-to-end smFRET analysis solution which also supports data filtering and graphing utilities. As the smFRET field is rapidly developing and diversifying, we designed FRETboard to grow with the needs of the user community; by allowing anyone to easily extend FRETboard with existing or future classification algorithms, our tool may continue to serve as a unifying web front-end for high-level users with both niche and all-purpose classification needs.

**Figure 1:**
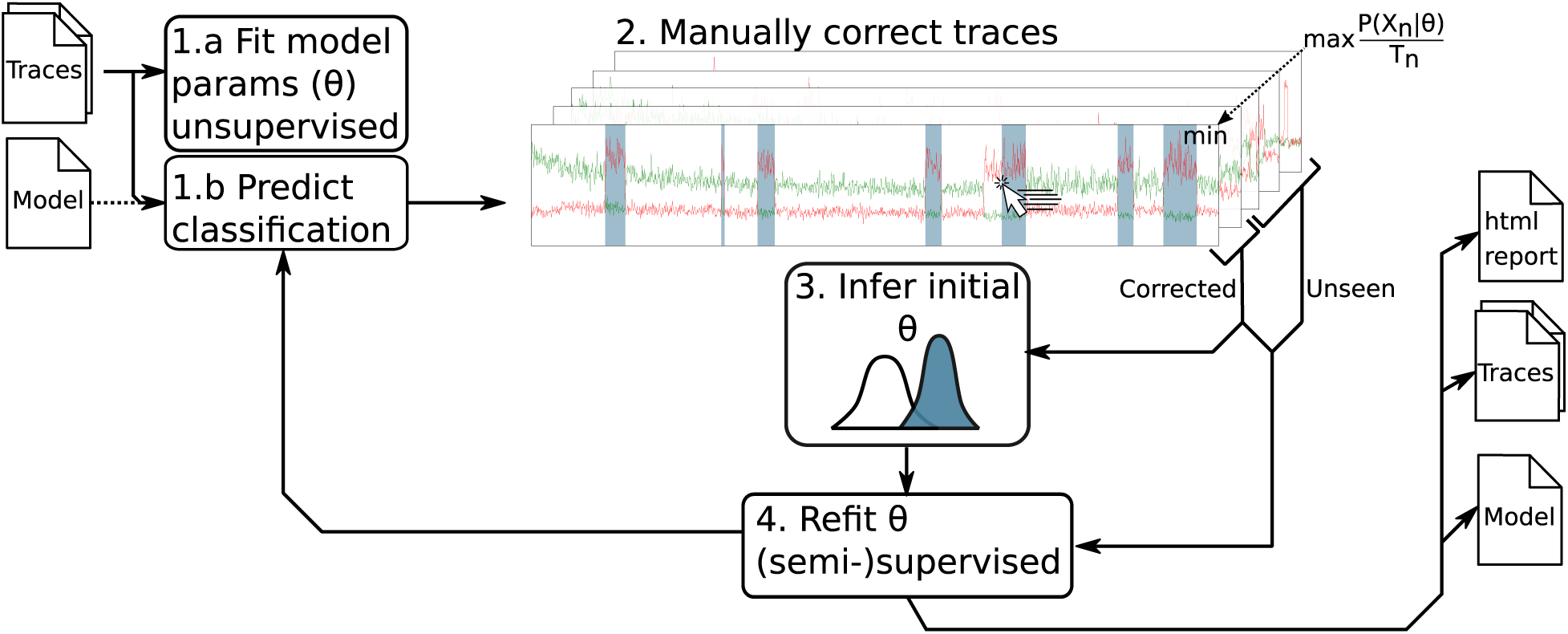
Our semi-supervised classification workflow for FRET trace classification, as implemented in FRETboard, is divided in 4 steps. (1a) FRET traces are uploaded to the web server, after which parameters (*θ*) of an initial model are fitted unsupervised and (1b) an initial trace classification is predicted. A suitable model generated during a previous FRETboard run can also be supplied to skip initial fitting. (2) The user is then shown the predicted classification of the trace at index *n, X*_*n*_, with duration *T*_*n*_, for which model fit was poorest based on duration-normalized trace probability given current model parameters (*P* (*X*_*n*_|*θ*)*/T*_*n*_), and is asked for manual correction. (3) The curated trace is then used to reinitialize the model, which (4) is then trained in a (semi-)supervised fashion. Steps 2 to 4 may repeated until model fit is deemed satisfactory by the user.

## 2 Analysis method

The FRET trace analysis method presented here is different from previous methods in three respects: the semi-supervised training approach, the structure of the models that are fitted and the features on which they were trained. Here we explain and justify our choices in each of these respects.

### 2.1 Semi-supervised model fitting

Briefly, an HMM fitting procedure needs to estimate a transition matrix, which contains the transition probabilities between each state, and one feature probability distribution for each state. Existing HMM-based FRET classification methods use an implementation of expectation maximization (EM) to alternate classification of the data and fitting the transition matrix and the probability distributions to the data, until parameters no longer change.

We address two notable downsides of unsupervised fitting in the context of FRET trace analysis. First, initial guesses are required for transition and emission parameters. Without prior knowledge these can be randomly generated, however their choice may exert significant influence on the eventual model parameters. Unlucky initialization may result in reaching a poor, local optimum, which becomes increasingly likely as the signal to noise ratio decreases. Repeated initialization and selection of the model with highest likelihood may mitigate this issue, but increases computational overhead. Second, the most likely state sequence after parameter optimization may not correspond well to classification by a trained human eye; limited data may cause an unsupervised fitting procedure to lump states that an expert would consider indicative of distinct molecular configurations, or inversely, split a state corresponding to a single configuration. In this case the parameter combination that would reproduce the expert labeling simply attains a different (i.e. not necessarily higher) optimum in the likelihood landscape.

Semi-supervised training offers a way to mitigate both issues. Initial emission distributions and transition probabilities can be estimated based on a manually labeled subset of all traces. Then training can proceed using the EM procedure on labeled and unlabeled data. The labels of classified examples need not be re-estimated and continue to steer the training procedure towards finding a model that reproduces the classification in the labeled traces, while still being able to fit patterns that were only present in the unlabeled traces.

We further improve on this analysis design by choosing to supervise traces to which the model assigned the lowest duration-normalized probability of the sequence. As these traces are the least well described by the model, it is more likely that the user’s guidance here steers the model towards a better fit. The fitting procedure is summarized in Figure 1, and more formally described in appendix C.

### 2.2 Model structures

We evaluate the performance of two HMM structures in a semi-supervised learning setting (Figure 2). Each structure implements transitions and/or emissions differently and can be extended to an arbitrary number of states. The “vanilla” structure produces a straight-forward fully-connected HMM sporting no further modifications. The “GMM-HMM” structure models emissions using a Gaussian mixture model (GMM) and adds additional states between neighboring states, which capture the distribution of features at the boundaries between transitions. The number of Gaussians per GMM is determined per state using a Bayesian information criterion (BIC)-selection procedure.

**Figure 2:**
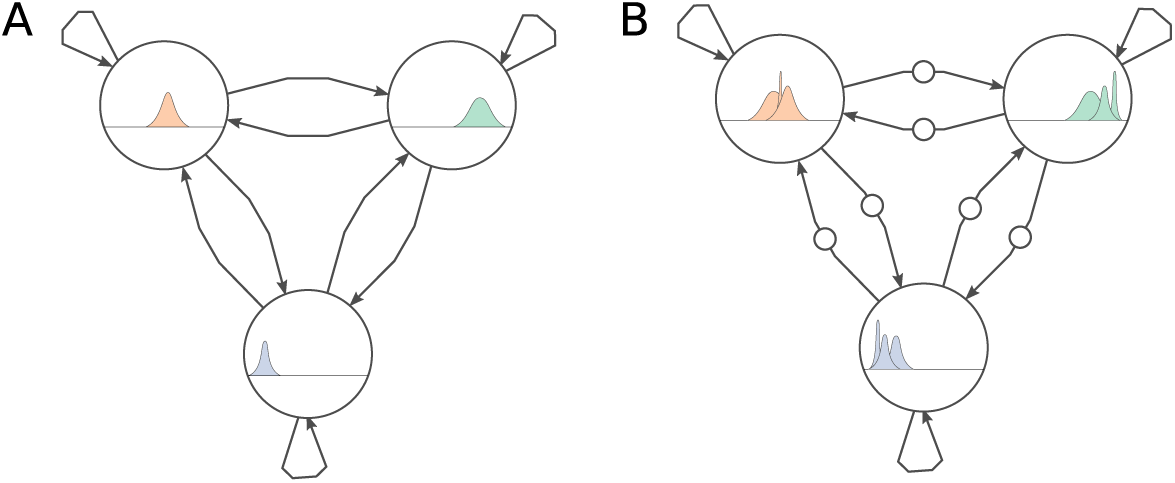
Hidden Markov model graph structures used in this work: (**A**) the plain “vanilla” structure, and (**B**) the “GMM-HMM” structure, in which each state contains a Gaussian mixture distribution and additional states for feature distributions at state transitions. Circles denote states with a characteristic feature distribution, edges denote transitions. Three-state structures are shown here, however similar structures with an arbitrary number of states can be constructed.

### 2.3 Features

We trained our models on a combination of four features. The proximity ratio *E*_*PR*_ is included as an approximation of FRET efficiency and is defined as:

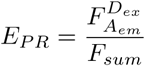

Here 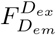 and 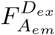 are the original donor and acceptor emission intensities. We also include the summed intensity 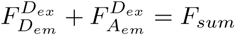 as a separate feature, as it is expected to aid in the detection of bleaching or absence of both dyes.

Furthermore we used two time-aggregated features that capture the variability of features over a sliding-window of five measurements; the Pearson correlation coefficient between 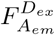 and 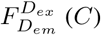 and standard deviation of *F*_*sum*_ 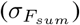. These features may aid models in capturing feature distributions characteristic for state transitions (Figure 2B). We specifically refrain from using 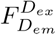 and 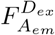 as features, as systematic variations frequently occur between experiments or even within the same experiment, which decreases the generalizability re-usability of a trained model.

### 2.4 Implementation

To facilitate application of our method we developed FRETboard, a browser-based graphical user interface (GUI) for semi-supervised training of segmentation and classification algorithms (Figure S1). In addition to intuitive example supervision, FRETboard offers users the flexibility to choose between model structures and opt which features to include. As we foresee that more suitable supervise-able classifiers may be proposed for the growing number of labeling schemes in the future, we also offer users the option to write custom algorithms and train them through the same FRETboard front-end. However due to the security risk of code injection that is inherent to running such custom code, users are advised to only allow this option on private machines that are not exposed to the public network.

Traces may be loaded in plain text, binary 64-bits or photon-HDF5 [13]-format. Given alternating laser excitation (ALEX) data, FRETboard can also correct optical data for fluorescent dye cross-talk, acceptor direct excitation, detection efficiencies and background emission (appendix D).

After the training procedure, the user may generate a report detailing feature distributions per state and transition rates. Transition rates are derived by taking transition probabilities from the fitted HMMs transition matrix (*A*), converting from discrete to continuous rates and multiplying by the frame rate *f*_*s*_, thus arriving at corrected transition rates *F* (Equation 1) [9].

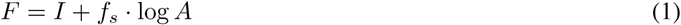

Here *I* is the identity matrix and log denotes the natural matrix logarithm operation. 95% confidence intervals (CIs) of transition rates are estimated by repeatedly extracting transition rates from bootstrapped data. The CI is then reported using the bootstrap standard deviation on each parameter. Note that bootstrapping CIs is applicable to almost any (semi-)supervised model, thus any user-defined algorithm can make use of the same method.

FRETboard is available as a web tool (https://www.bioinformatics.nl/FRETboard), thus freeing users from the burden of installation and maintenance, but can also be used and hosted on a private server. FRETboard was written in python 3.7 (https://www.python.org). The GUI was implemented using the Bokeh interactive visualization library (v1.4.0)[14] (Fig. S1). Included HMM model structures were implemented using pomegranate (v0.13.4) [15] and scikit-learn (v0.21.2) [16].

## 3 Data analysis and performance evaluation

Below we validate our analysis method on four *in silico* and four *in vitro* data sets. To demonstrate the flexibility of our method, the different sets were simulated or recorded assuming a variety of realistic labeling schemes. All data sets used here are freely available (https://git.wageningenur.nl/lanno001/fretboard_data). All FRETboard runs were performed on a laptop running Ubuntu 18.04, on four CPU cores (Core i7 1.80GHz, Intel Corp.) with 4GB of memory. In total we supervised ten traces for each data set (3% and 10% of the total number of reads for *in silico* and *in vitro* data sets respectively), making use of the four described features (*E*_*PR*_, *F*_*sum*_, *C*, and 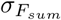).

We assessed how well semi-supervised HMMs were able to reproduce ground truth parameters derived from the simulated state sequences or, in the case of *in vitro* data, from the manual labeling. To test whether predicted *E*_*PR*_ distributions attain a mean comparable to the ground truth, we apply the two one-sided *t*-tests (TOST) procedure [17]. That is, for a given state *s* the predicted mean of 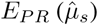 and ground truth mean (*µ*_*s*_) are calculated and two one-sided *t*-tests are employed to test 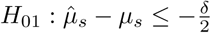 and 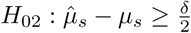 versus 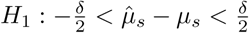. A rejection of both hypotheses implies that the difference between emission means is significantly smaller than *δ E*_*PR*_ percentage points. Here we test for a maximum deviation of five or ten percentage points (*δ* = 0.05 and *δ* = 0.1 respectively), which we consider sufficiently accurate for many current applications applications. We report the TOST *p*-value for a given *δ* = *δ\** as *p*_*δ*=*δ\**_. Reported 95% CIs around estimated transition rates were calculated using FRETboard’s built-in bootstrapping method, using a bootstrap size of 100.

### 3.1 Performance on *in silico* data

To demonstrate the flexibility of our approach, we simulated FRET traces based on three different labeling schemes using a Markov Chain Monte Carlo process to generate state sequences with added Gaussian noise and classified them using a semi-supervised vanilla HMM. Briefly, the first two data sets contain two and three FRET states respectively, which are separable based on *E*_*PR*_ only (Figure S2A, B). The third data set contains three states, of which the third is identical to the second in its proximity ratio but has a different transition rate, making it a ‘degenerate state’ (Figure S2C). For a full description of the simulation methodology see appendix A.

In all cases, estimated mean *E*_*PR*_ significantly differed less than 5 percentage points from the ground truth mean (*p*_*δ*=0.05_ << 0.001) (Figure 3 A-C). Most ground truth transition rates fell well within bootstrapped 95%-CIs around the predicted rates (Figure 3 E-G). If a degenerate states was present (Figure 3 C, G), transition rate estimates deviated somewhat from the ground truth, indicating that a model structure that is better equipped to detect these is in order for such cases [e.g. 11]. Apart from the manual curation, no further parameter tuning or other user input was required, demonstrating that semi-supervised training provides the expected flexibility while maintaining accuracy.

**Figure 3:**
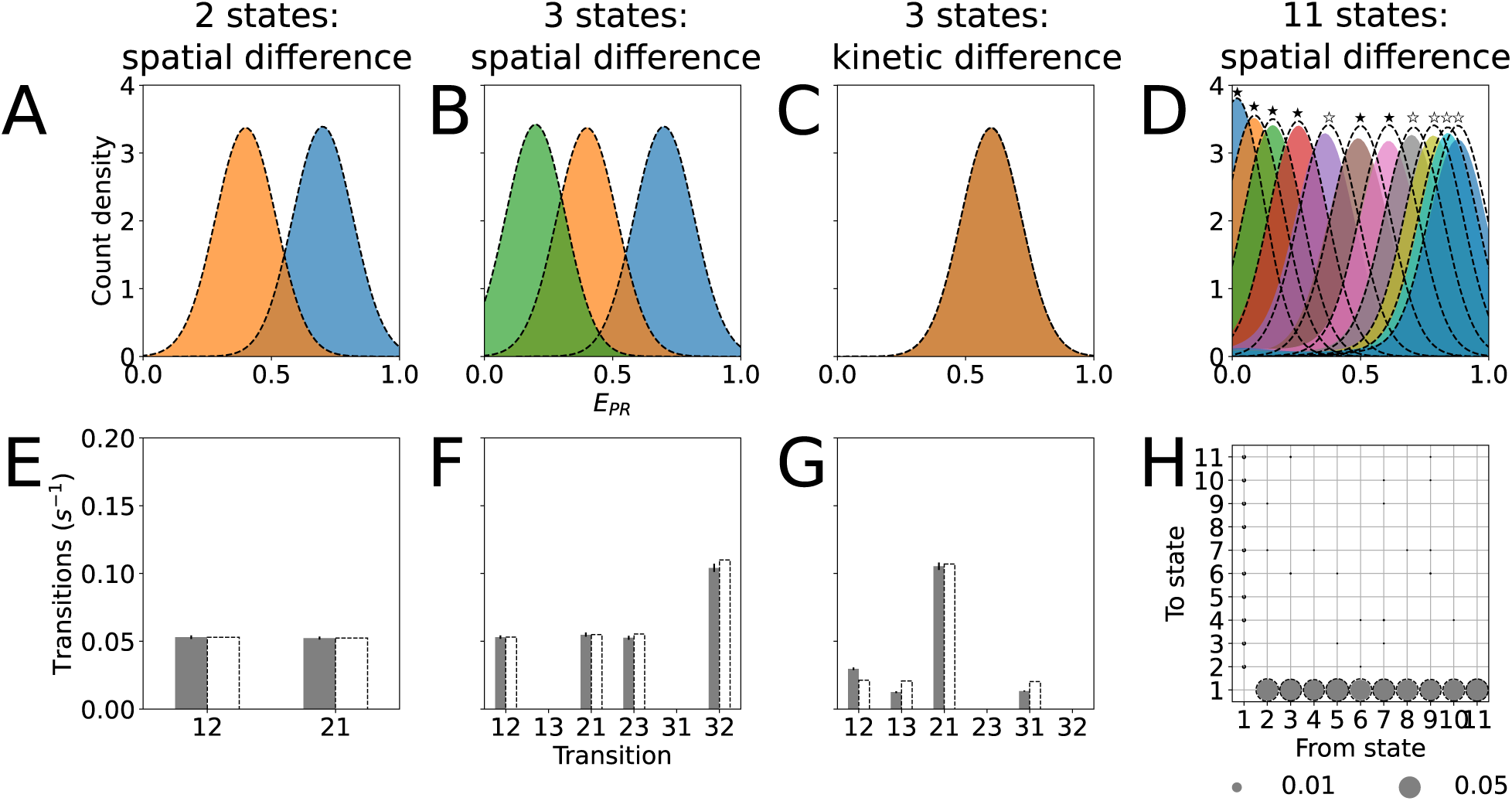
kernel density estimates of *E*_*PR*_ distributions per state (**A-D**) and transition rates (**E-H**) as estimated by a semi-supervised hidden Markov model on four simulated data sets based on different labeling schemes; (**A**,**E**) producing two types of spatially separable FRET events, i.e. with different proximity ratios (*E*_*PR*_), (**B**,**F**) three spatially separable states, (**C**,**G**) three states of which one containing no donor or acceptor signal and the others exhibiting only a kinetic difference, i.e. separable by transition rate and (**D, H**) eleven spatially separable states. Ground truth parameters are displayed as dotted lines in each figure. In D, symbols above *E*_*PR*_ distributions denote how much estimated means significantly differ from the ground truth at most (⋆: 0.05, ✶: 0.1). Solid black lines in E-G indicate 95% bootstrapped CIs. Circle surface in H indicates transition rate.

To stress-test our method on a more difficult case, we generated a data set in which eleven FRET states of differing *E*_*PR*_-levels were present (Figure S2D). The mean *E*_*PR*_ values were distributed such that their corresponding donor-acceptor distances were evenly distributed, thus causing lower and higher ends of the *E*_*PR*_ spectrum to be more densely crowded with states. Additionally, donor and acceptor signals were generated using mixtures of Gaussians to emulate noise in less ideal measuring circumstances.

During training, it became clear that the vanilla model structure lacks the complexity to properly fit this data set, as each trace presented after a new round of training continued to require correction by the user without further increasing training accuracy. Indeed, transition rates and *E*_*PR*_ distributions estimated using the vanilla structure differ significantly from ground truth values (Figure S3). We thus continued our analysis using the GMM-HMM structure. Six of eleven *E*_*PR*_ distribution means significantly differed less than five percentage points from the ground truth means (*p*_*δ*=0.05_ << 0.001, Figure 3D), with the remaining five differing less than ten percentage points (*p*_*δ*=0.1_ << 0.001) and transition rate estimates were closer to ground truth values than vanilla HMM estimates (3H). This demonstrates another strength of our method; as the user discovers that simpler models are not delivering the desired accuracy, the training procedure is light enough that a more elaborate model can be selected on the fly, after which the analysis can continue without extra effort.

### 3.2 Performance on *in vitro* data

We further validated our method on experimental data generated under immobilization schemes often used in single-molecule FRET, each marked by different classification challenges. Similar to our simulations, our *in vitro* data contains up to two types of FRET events, which may be discernible by proximity ratio or transition rate. Lacking knowledge of the state sequence, we manually classified our data sets and refer to this classification as the ground truth. A more extensive description of experimental methods can be found in appendix B. All experimental data was analyzed using the GMM-HMM model structure, as the vanilla structure did not show a satisfactory increase in classification quality as training progressed (Figure S4).

First we designed an experiment in which a donor (Cy3)-labeled single-stranded (ss) DNA, containing a target site A, is immobilized through biotin-streptavidin conjugation on a quartz slide (Figure 4A). Upon binding of an acceptor (Cy5)-labeled eight-nucleotide ‘imager’ strand to site A, FRET events marked by anti-correlated donor and acceptor signals are produced (Figure 4D). Similar labeling schemes have previously seen application in point accumulation for imaging in nano-scale topography (PAINT) methods and the study of on- and off-rates (*k*_*on*_ and *k*_*off*_) in biological systems [18, 19]. In this labeling scheme, bleaching of the donor dye due to continuous excitation occurs frequently, which may negatively impact kinetics analysis. Instead of requiring the user to laboriously remove bleaching events, we capture them in a separate state while training our HMM. This bleached state may then be discarded prior to further analysis. Following this approach, we find that ground truth transition rates for ground to high-FRET state and vice versa indeed fall within their respective estimated CIs (0.119*s*^−1^ versus *CI* : (0.095 − 0.122) and 0.546*s*^−1^ versus *CI* : (0.468 − 0.578) respectively, Figure 4M). Estimated *E*_*PR*_ values (0.151 and 0.816 for ground and high-FRET states respectively) significantly differ by less than five percentage points from ground truth values *p*_*δ<*0.05_ << 0.01, Figure 4I).

**Figure 4:**
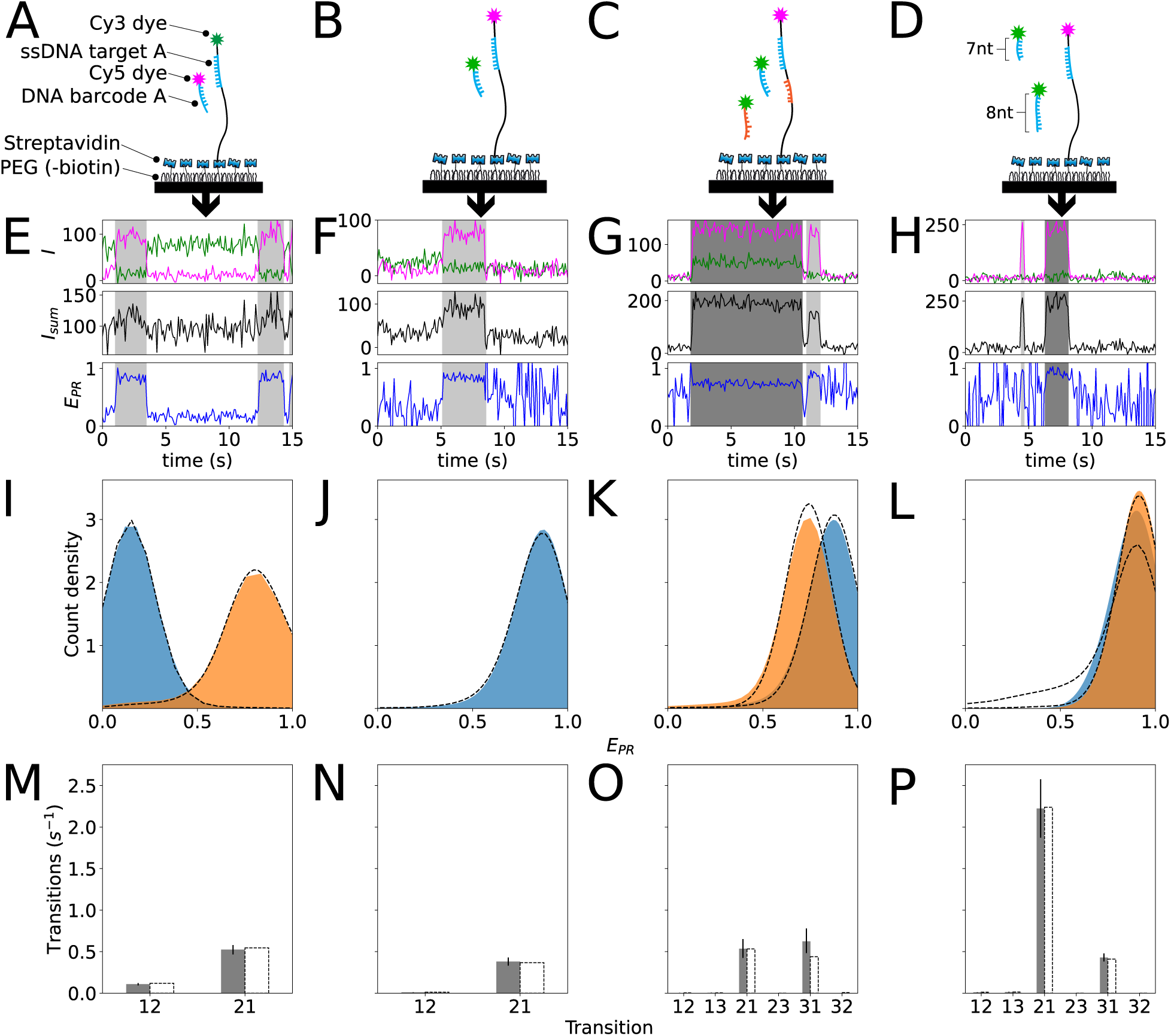
Used labeling schemes (**A-D**), examples of events produced as correctly found by semi-supervised HMM classification (**E-H**), kernel density estimates of *E*_*PR*_ distributions per state (**I-L**) and estimated transition rates (**M-P**) for four different labeling schemes on which our semi-supervised HMM fitting method was evaluated. From left to right, these labeling schemes were producing single-type FRET events using an immobilized donor (**A**) or an immobilized acceptor (**B**), two types of events producing high- and mid-FRET events (**C**) and two types of kinetically different events (**D**). In J-L non-FRET state is omitted as *E*_*PR*_ values are meaningless for labeling schemes in which the donor is not immobilized. In I-P, dotted lines represent ground truth values based on manual labeling and in M-P solid lines denote bootstrapped 95% CIs.

We also performed the reverse experiment, in which the acceptor is immobilized and the donor is attached to the imager strand (Figure 4B). Although this labeling scheme does not suffer from dye bleaching as much, the lack of anti-correlation in the signal is expected to increase the difficulty of classification. Nonetheless, here too the predicted *E*_*PR*_ distribution mean of the high-FRET state (0.844) differed from the ground truth value by less than 5 percentage points (*p*_*δ*=0.05_ << 0.01, Figure 4J). As no dye is observable in the ground state under this labeling scheme, its *E*_*PR*_ value is meaningless and not analyzed here. The ground truth transition rate from high-FRET to ground state fell within its predicted 95% CI (0.367*s*^−1^ versus *CI* : (0.333 − 0.429)), while the rate for ground to high-FRET state was slightly underestimated (0.012*s*^−1^ versus *CI* : (0.007 − 0.011), Figure 4N).

Next, we evaluated performance in two scenarios where two FRET states are present. For these experiments we followed the same experimental procedure, but simultaneously flushed in two types of donor-bound free-floating imager strands, at a 1:1 ratio (Figure 4D,E).

In the first experiment, the second imager strand was complementary to a second target site B at 15nt from the acceptor – 10nt further than target site A – where imager strand binding should produce an intermediate *E*_*PR*_ (Figure 4F). Upon analysis, the GMM-HMM model found *E*_*PR*_ means of 0.85 and 0.72 for states 2 and 3 respectively, matching the ground truth state means (*p*_*δ<*0.05_ *<* 0.01, Figure 4J). Most transition rate estimates fell within the predicted 95% CIs, except for that from mid-FRET (3) to ground state (1)(0.439*s*^−1^ versus *CI* : (0.483 − 0.780)). Upon inspection of traces, we found that several short mid-FRET events had been erroneously detected in noisy ground state stretches – a common occurrence in smFRET analysis and therefore not explicitly accounted for by e.g. removing traces from analysis manually.

In the second experiment, site A was targeted with a second imager strand of 7nt – 1nt shorter than its counterpart – which should increase the off-rate and produce a degenerate state (Figure 4G). Here too our GMM-HMM produced parameter estimates close to ground truth values on traces containing degenerate states, which is surprising given our results on *in silico* data. Predicted transition rates from state 2 to ground state were higher at 2.22*s*^−1^(*CI* : 1.87 − 2.57), than that of state 3 – 0.43*s*^−1^(*CI* : 0.382 − 0.479) –, which resembled transition rates seen in other *in vitro* experiments (Figure 4O). Presumably, state 2 corresponds to the annealing of the shorter 7nt imager strand. Both are in close agreement with ground truth rates (2.24*s*^−1^ and 0.411*s*^−1^ respectively).

## 4 Discussion

We show that semi-supervised classification models, in particular hidden Markov models (HMMs), are capable of capturing properties of FRET events in a wide array of realistic experimental scenarios, using a combination of input features derived from the original donor and acceptor dye emission intensities. We also provide an HMM structure that is overall better suited for semi-supervised learning than the straight-forward fully connected model, and provide a particular advantage in noisy real-world and complex data sets containing more than two states.

An important caveat of our performance assessment – and supervised learning in general – is that the user is assumed to know the number of states to fit and what characteristics to look for in manual labeling. With our method we give the responsibility of proper classification to the user, embracing the pros and cons of user input; on the one hand, it allows for efficient training and yields results that match the user’s intuition, on the other hand it matches mistakes that the user may make. Such mistakes could occur for instance in the classification of traces after dyes have bleached. If the user is only interested in the average FRET efficiency of non-ground state events, it may make sense to classify bleached stretches as ground state – a course of action for which the GMM-HMM structure in particular lends itself due to its flexibility. However, if kinetic rate estimates are of importance, results are more accurate if bleaching is assigned a state of its own which can then be removed prior to analysis, as was done in the analysis presented here.

To accommodate for the intensive user interaction required for this method we developed FRETboard, an intuitive browser-based tool that allows data preparation, model training, classification and report generation. Importantly, FRETboard can easily be extended to other model structures than the ones presented in this work; as many superviseable HMM flavors and entirely different classifiers exist and may be a better fit than the models currently included for certain experimental data, we encourage users to design their own classifiers and train them through the FRETboard interface. In consultation with the authors of such custom classifiers, these may also be included in future releases of FRETboard. This would allow it to become a unifying front end for FRET trace analysis, with back end support for the expanding variety of smFRET experimental methods.

## Appendices

### A Trace simulation

Simulated traces were generated by using a custom code written in Matlab (R2019b, The MathWorks, Inc.). A Markov chain Monte-Carlo (MCMC) procedure was used to generate hidden system state paths according to a given transition matrix. The donor and acceptor fluorescence intensity traces were then generated from the hidden paths, using given emission distributions of single or mixed Gaussian per state. The widths of the Gaussian distributions were set to the square root of the mean to mimic the photon shot noise contribution. Additional random background noise was further added to simulate the dark count of the photo detector and the electronics shot noise. The traces were then downsampled to one tenth of the original length to simulate discrete measurement behaviour on a continuous biological process. For each dye labeling scheme, 300 single molecule traces were generated and used to test the performance of FRETboard. The length of each individual trace was randomly chosen between 12000 to 18000 data points before downsampling in all except the 11-state case, for which 24000 to 36000 data points were used. The simulation code is freely available upon request.

### B Extended experimental methods

#### B.1 Single-molecule setup

All experiments were performed on a custom-built microscope setup. An inverted microscope (IX73, Olympus) with prism-based total internal reflection is used. In combination with a 532 nm DPSS laser (Compass 215M/50mW, Coherent). A 60x water immersion objective (UPLSAPO60XW, Olympus) was used for the collection of photons from the Cy3 and Cy5 dyes on the surface, after which a 532 nm long pass filter (LDP01-532RU-25, Semrock) blocks the excitation light. A dichroic mirror (635 dcxr, Chroma) separates the fluorescence signal which is then projected onto an EM-CCD camera (iXon Ultra, DU-897U-CS0-#BV, Andor Technology). A series of EM-CDD images was recorded using custom-made program in Visual C++ (Microsoft). Time traces were extracted from the EM-CDD images using IDL (ITT Visual Information Solution).

#### B.2 Single-molecule data acquisition

To avoid non-specific binding of DNA to the surface, quartz slides were PEGylated as previously described [20]. Briefly, acidic piranha etched quartz slides (Finkenbeiner) were passivated twice with polyethylene glycol (PEG). The first round PEGylation was performed with mPEG-SVA (Laysan) and PEG-biotin (Laysan), followed by a second round of PEGylation with MS(PEG)4 (ThermoFisher). After assembly of a microfluidic chamber, the slides were incubated with 20 *µ*L streptavidin (0.1 mg/mL, ThermoFisher) for 2 minutes. Excess streptavidin was removed with 100 *µ*L T50. Next, for single-molecule experiments we immobilized 50 *µ*L of 100 pM Cy5 (or Cy3) labelled target DNA for 2 minutes, unbound DNA was washed with 100 *µ*L T50, followed by 100 *µ*L of buffer A (50 mM Tris-HCl, pH 8.0, 500 mM NaCl). Next, we injected 50 *µ*L of 5 nM imager strands in imaging buffer (50 mM Tris-HCl, pH 8.0, 500 mM NaCl, 0.8% glucose, 0.5 mg/mL glucose oxidase (Sigma), 85 *µ*g/mL catalase (Merck) and 1 mM Trolox (Sigma)). The single-molecule FRET experiments were performed at room temperature (23 *±* 2 degC).

### C Semi-supervised model fitting

If manually labeled traces are available, the set of all parameters (*θ*) for a Hidden Markov model (HMM) can be fitted in a supervised fashion; emission probabilities may be estimated by fitting a probability distribution to features for each state, while transition probabilities are deduced from the state sequence. To fit Gaussian distributions to our features for each state *i* given labeled traces *X*_*n*_, *n* = 1*…N*, of variable length *T*_*n*_, we calculate the mean 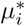 and standard deviation 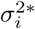 over all measurements *x*_*nt*_ of which the corresponding label *L*_*nt*_ equals *i*:

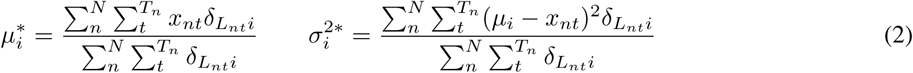

Here 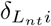 equals 1 if *L*_*nt*_ = *i*, and 0 if *L*_*nt*_ ≠ *i*. The transition probabilities are estimated as the ratio of transitions between two states over the number of occurrences of the departure state:

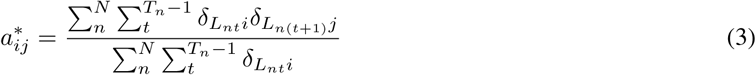

Here 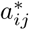 denotes the estimated entry (*i, j*) in the HMM’s transition matrix, i.e. the probability of transitioning from state *i* to *j*. Finally, the starting probabilities per state can be derived by taking the fraction of labeled traces that start with a given state:

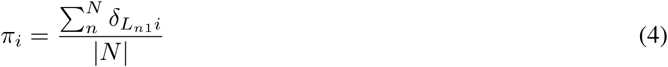

Manually labeling sufficient data to capture all distinctive patterns in traces is laborious, thus HMMs are often fitted unsupervised, using an implementation of expectation maximization known as Baum-Welch training. By iteratively alternating between re-estimating model parameters and producing the most likely labeling given model parameters and data, an HMM can be fitted without prior knowledge of either the correct sequence labeling or model parameters. In that case the contribution of each sample to the parameters for the emission distribution of a class *i* and a given transition *a*_*ij*_ is weighted by the probabilities of assignment to that class *γ*_*i*_(*n, t*) and the transition occurring *ξ*_*ij*_(*n, t*) respectively, given the current model parameters:

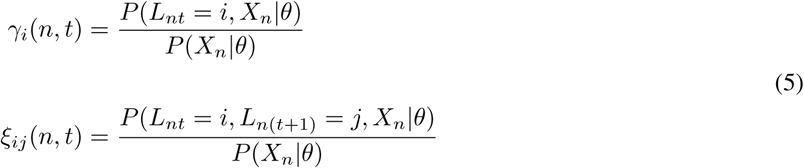

Fitting using the supervised procedure effectively sets the probabilities for observed states and transitions to 1 and all others to 0 by means of 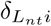 in equations 2 and 3.

As noted in the main text, Baum-Welch training requires initial values for transition and emission parameters, and given its greedy nature may not converge to a satisfactory optimum. If semi-supervised learning is employed, initial emission distributions and transition probabilities can be estimated using a manually labeled subset of all traces and equations 2 and 3. Then training can proceed using the Baum-Welch procedure on labeled and unlabeled data, where the labels of classified examples need not be re-estimated (equations 6). In this way, the training procedure may be steered towards finding a model that reproduces the classification in the labeled traces, while still being able to fit patterns that were only present in the unlabeled traces.

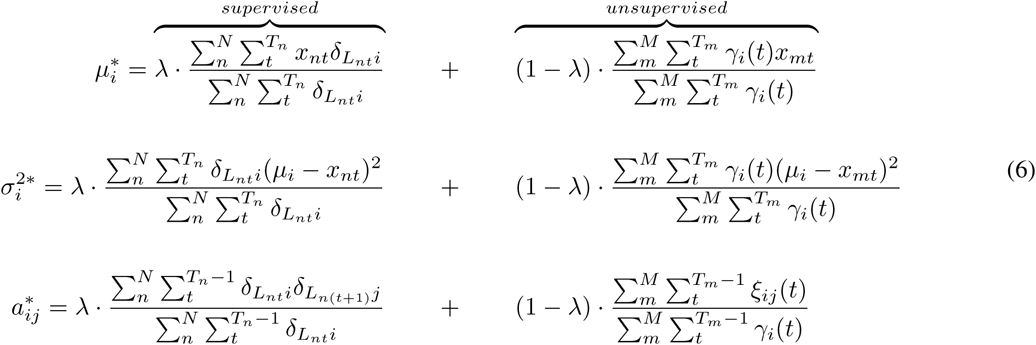

here *N* denotes the number of manually labeled traces and *M* denotes the number of unlabeled traces. We may choose to increase the additional tuning parameter *λ* if we find that the model does not reproduce our labeled sequences sufficiently well anymore. Note that for *λ* = 1 the procedure resorts to fully supervised fitting, which requires no Baum-Welch training. By default, *λ* is set to 0.5, meaning that all supervised and unsupervised traces carry the same weight.

### D Trace filtering using FRETboard

As FRET efficiency is influenced by background irradiance, detector efficiency differences, cross-talk between acceptor and donor channels, direct excitation of acceptor dyes and background emission, it is advisable to apply some form of filtering to raw traces prior to classifier training.

FRETboard offers an automated per-trace background subtraction method for donor and acceptor channels separately. Using the DBSCAN clustering algorithm the lowest intensity level in the channel is detected and subtracted. Prerequisite for proper functioning of this adaptive filter is that the trace does contain a stretch of background-level intensity irradiance. Stringency of this filter is controlled by the tunable *ϵ* parameter, which is defined as the maximum distance between a cluster’s core point and distant points of the same cluster. A preset value of *ϵ* = 15 was found to work in all data presented here.

If alternating laser excitation (ALEX) data is available, FRETboard can automatically apply cross-talk, direct excitation and detector efficiency corrections as described in [21], by marking certain states as *D*(onor)-only and *A*(cceptor)-only. FRETboard will subsequently perform the necessary calculations to estimate and apply correction coefficients.

### E Supplementary figures

**Figure S1:**
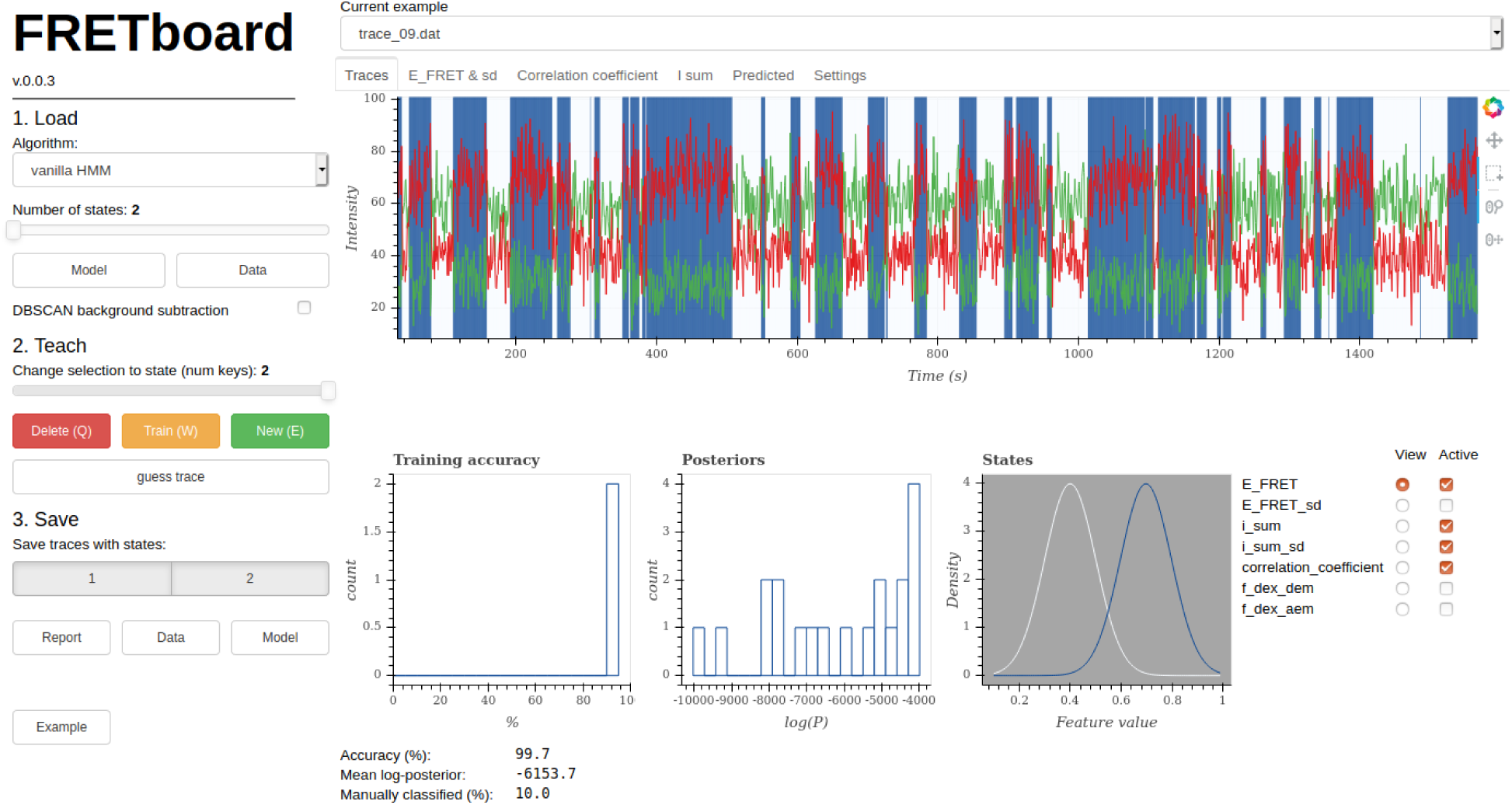
Controls in the graphical user interface are divided in three steps: (1) loading data and model structure, (2) iteratively teaching the chosen model structure to recognize events as the user sees fit and (3) saving a report with statistics and publication-ready figures, the raw classified data for further processing and the model for future use on similar datasets.

**Figure S2:**
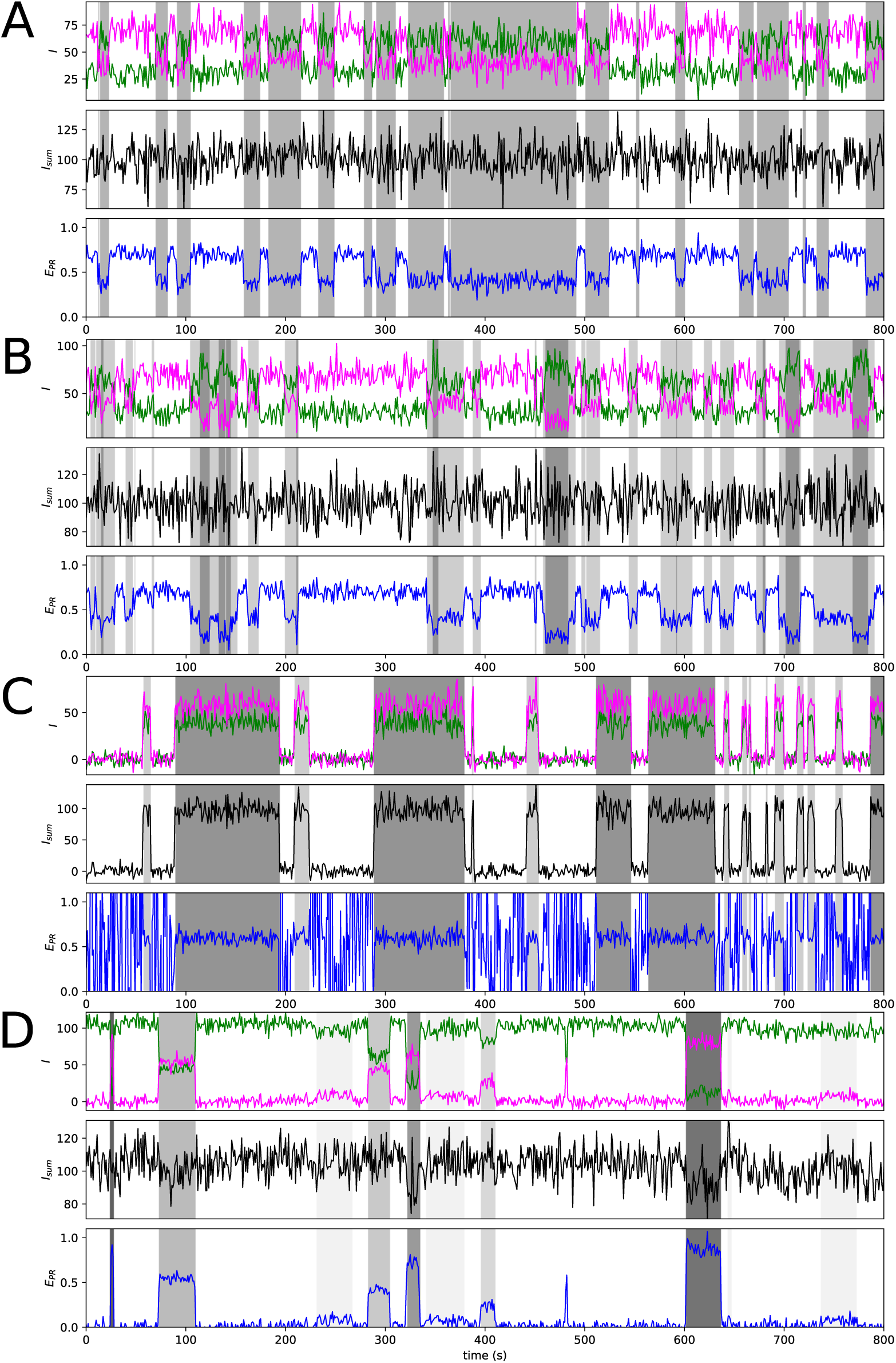
Example sections of simulated FRET traces and ground truth classification for four scenarios: **(A)** one FRET state, **(B)** one FRET state with immobilized acceptor and mobile donor **(C)** two FRET states and **(D)** ten FRET states. White background denotes the ground state while different shades of gray denote different states.

**Figure S3:**
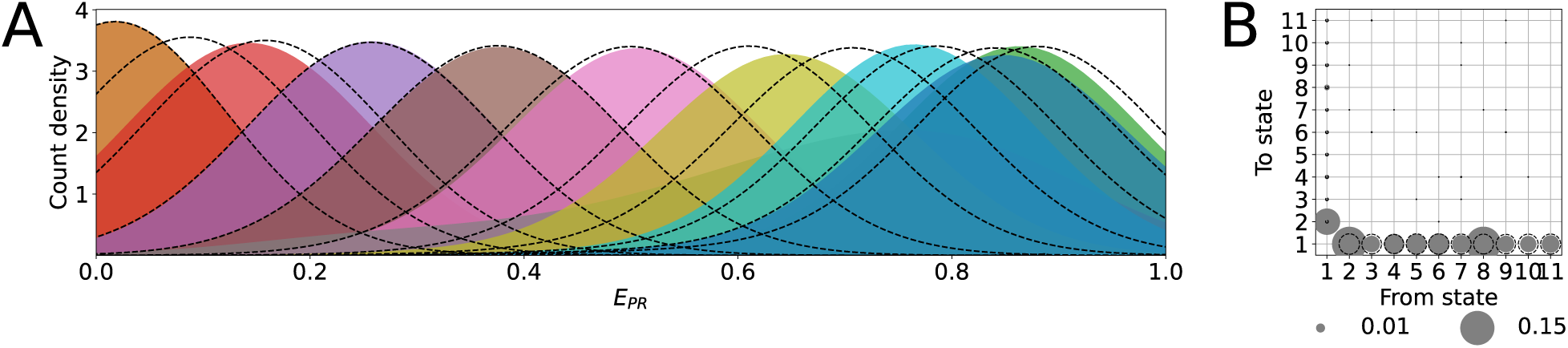
Kernel density estimates of *E*_*PR*_ distributions per state (**A**) and transition rates (**B**) as estimated by a semi-supervised vanilla hidden Markov model on a simulated dataset containing eleven separable states. Ground truth parameters are displayed as dotted lines in each figure. In B, circle surface indicates transition rate. In comparison, analysis using the GMM-HMM model structure (Figure 3D,H) returned parameter estimates closer to ground truth values for both transition rates and *E*_*PR*_ distributions per state.

**Figure S4:**
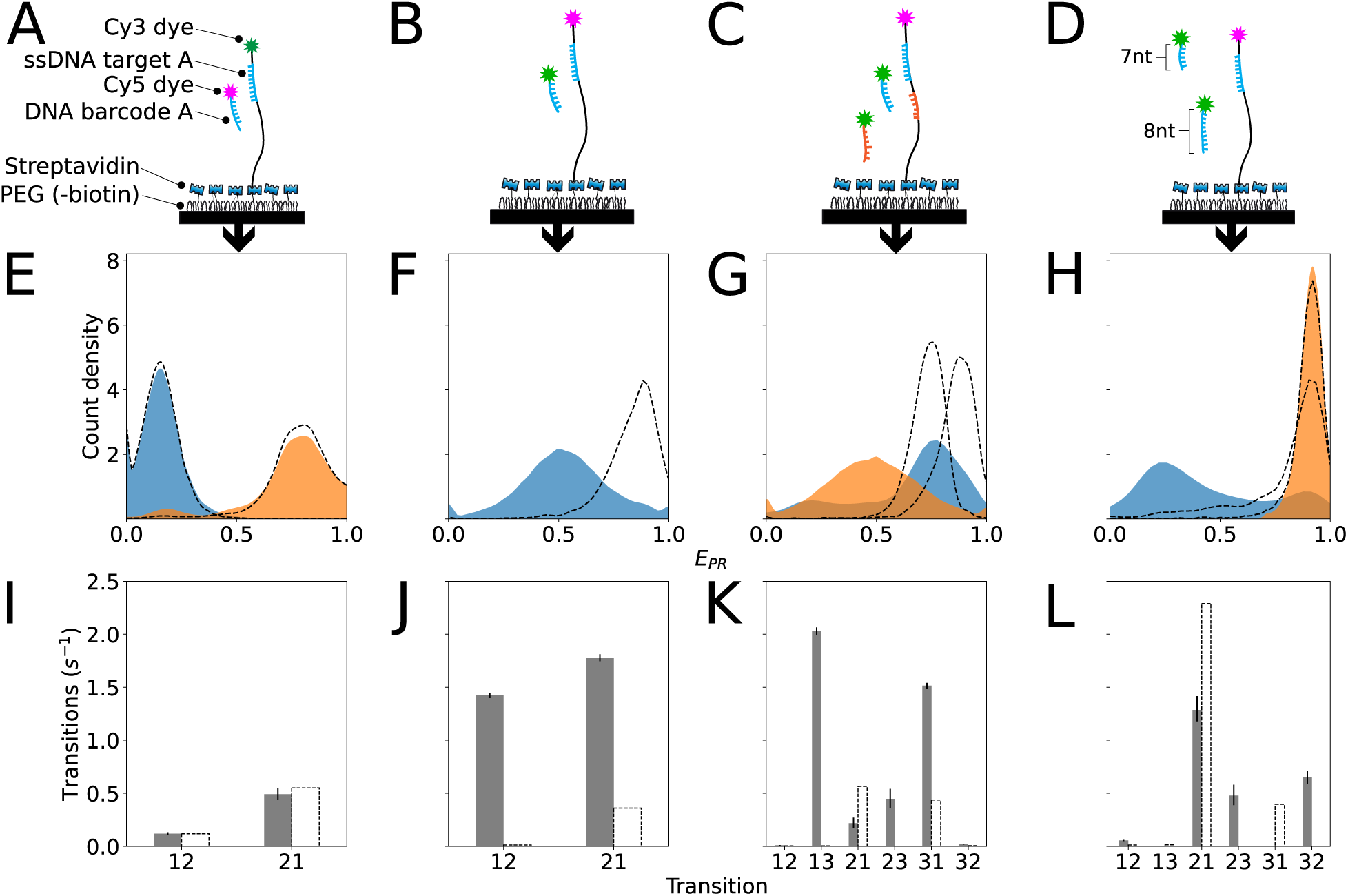
Used labeling schemes (**A-D**), kernel density estimates of *E*_*PR*_ distributions per state (**E-H**) and estimated transition rates (**I-L**) for four different labeling schemes on which our semi-supervised HMM fitting method with vanilla structure was evaluated. From left to right, these labeling schemes were producing single-type FRET events using an immobilized donor (**A**) or an immobilized acceptor (**B**), two types of events producing high- and mid-FRET events (**C**)and two types of kinetically different events (**D**). In E-L, dotted lines represent ground truth values based on manual labeling and in M-P solid lines denote bootstrapped 95% CIs. In comparison, analysis using the GMM-HMM model structure (Figure 4I-P) returned parameter estimates closer to ground truth values for both transition rates and *E*_*PR*_ distributions per state.

## References

[1] Achillefs N Kapanidis, Emmanuel Margeat, Sam On Ho, Ekaterine Kortkhonjia, Shimon Weiss, and Richard H Ebright. Initial transcription by rna polymerase proceeds through a dna-scrunching mechanism. Science, 314(5802):1144–1147, 2006.

[2] Allan Chris M Ferreon, Yann Gambin, Edward A Lemke, and Ashok A Deniz. Interplay of *α*-synuclein binding and conformational switching probed by single-molecule fluorescence. Proc. Natl. Acad. Sci. U. S. A., 106(14):5645–5650, 2009.

[3] Michael Börsch and Thomas M. Duncan. Spotlighting motors and controls of single FoF1-ATP synthase. Biochem. Soc. Trans., 41(5):1219–1226, 2013.

[4] Eitan Lerner, Antonino Ingargiola, and Shimon Weiss. Characterizing highly dynamic conformational states: The transcription bubble in RNAP-promoter open complex as an example. J. Chem. Phys., 148(12):123315, 2018.

[5] Matthew D Newton, Benjamin J Taylor, Rosalie PC Driessen, Leonie Roos, Nevena Cvetesic, Shenaz Allyjaun, Boris Lenhard, Maria Emanuela Cuomo, and David S Rueda. DNA stretching induces Cas9 off-target activity. Nat. Struct. Mol. Biol., 26(3):185–192, 2019.

[6] Viktorija Globyte, Seung Hwan Lee, Taegeun Bae, Jin-Soo Kim, and Chirlmin Joo. CRISPR/Cas9 searches for a protospacer adjacent motif by lateral diffusion. EMBO J., 38(4):e99466, 2019.

[7] Stephan Uphoff, Seamus J Holden, Ludovic Le Reste, Javier Periz, Sebastian Van De Linde, Mike Heilemann, and Achillefs N Kapanidis. Monitoring multiple distances within a single molecule using switchable FRET. Nature Methods, 7(10):831, 2010.

[8] Sean A. McKinney, Chirlmin Joo, and Taekjip Ha. Analysis of single-molecule FRET trajectories using hidden Markov modeling. Biophys. J., 91(5):1941–1951, 2006.

[9] Max Greenfeld, Dmitri S. Pavlichin, Hideo Mabuchi, and Daniel Herschlag. Single Molecule Analysis Research Tool (SMART): an integrated approach for analyzing single molecule data. PLoS ONE, 7(2):30024, 2012.

[10] Jan Willem van de Meent, Jonathan E. Bronson, Chris H. Wiggins, and Ruben L. Gonzalez. Empirical Bayes methods enable advanced population-level analyses of single-molecule FRET experiments. Biophys. J., 106(6):1327–1337, 2014.

[11] Sonja Schmid, Markus Götz, and Thorsten Hugel. Single-molecule analysis beyond dwell times: demonstration and assessment in and out of equilibrium. Biophys. J., 111(7):1375–1384, 2016.

[12] Johannes Thomsen, Magnus B. Sletfjerding, Stefano Stella, Bijoya Paul, Simon Bo Jensen, Mette G. Malle, Guillermo Montoya, Troels C. Petersen, and Nikos S. Hatzakis. DeepFRET: Rapid and automated single molecule FRET data classification using deep learning. bioRxiv, 2020. doi: 10.1101/2020.06.26.173260.

[13] Antonino Ingargiola, Ted Laurence, Robert Boutelle, Shimon Weiss, and Xavier Michalet. Photon-HDF5: An open file format for timestamp-based single-molecule fluorescence experiments. Biophys. J., 110(1):26–33, 2016.

[14] Bokeh Development Team. Bokeh: Python library for interactive visualization, 2019.

[15] Jacob Schreiber. Pomegranate: fast and flexible probabilistic modeling in python. J. Mach. Learn. Res., 18(164):1–6, 2018.

[16] Fabian Pedregosa, Gael Varoquaux, Alexandre Gramfort, Vincent Michel, Bertrand Thirion, Olivier Grisel, Mathieu Blondel, Peter Prettenhofer, Ron Weiss, Vincent Dubourg, Jake Vanderplas, Alexandre Passos, David Cournapeau, Matthieu Brucher, Matthieu Perrot, and Édouard Duchesnay. Scikit-learn: Machine learning in Python. J. Mach. Learn. Res., 12:2825–2830, 2011.

[17] Donald J Schuirmann. A comparison of the two one-sided tests procedure and the power approach for assessing the equivalence of average bioavailability. J. Pharmacokinet. Biopharm., 15(6):657–680, 1987.

[18] Jongjin Lee, Sangjun Park, Wooyoung Kang, and Sungchul Hohng. Accelerated super-resolution imaging with FRET-PAINT. Mol. Brain, 10(1):63, 2017.

[19] Mike Filius, Tao Ju Cui, Adithya N. Ananth, Margreet W. Docter, Jorrit W. Hegge, John van der Oost, and Chirlmin Joo. High-speed super-resolution imaging using protein-assisted dna-paint. Nano Lett., 20(4):2264–2270, 2020.

[20] Stanley D Chandradoss, Nicole T Schirle, Malwina Szczepaniak, Ian J MacRae, and Chirlmin Joo. A dynamic search process underlies microrna targeting. Cell, 162(1):96–107, 2015.

[21] Nam Ki Lee, Achillefs N. Kapanidis, You Wang, Xavier Michalet, Jayanta Mukhopadhyay, Richard H. Ebright, and Shimon Weiss. Accurate FRET measurements within single diffusing biomolecules using alternating-laser excitation. Biophys. J., 88(4):2939–2953, 2005.

